# Structure and assembly of calcium homeostasis modulator proteins

**DOI:** 10.1101/857698

**Authors:** Johanna L Syrjanen, Kevin Michalski, Tsung-Han Chou, Timothy Grant, Shanlin Rao, Noriko Simorowski, Stephen J. Tucker, Nikolaus Grigorieff, Hiro Furukawa

## Abstract

Biological membranes of many tissues and organs contain large-pore channels designed to permeate a wide variety of ions and metabolites. Examples include connexin, innexin, and pannexin, which form gap junctions and/or *bona fide* cell surface channels. The most recently identified large-pore channels are the calcium homeostasis modulators (CALHMs), which permeate ions and ATP in a voltage-dependent manner to control neuronal excitability, taste signaling, and pathologies of depression and Alzheimer’s disease. Despite such critical biological roles, the structures and patterns of oligomeric assembly remain unclear. Here, we reveal the first structures of two CALHMs, CALHM1 and CALHM2, by single particle cryo-electron microscopy, which show novel assembly of the four transmembrane helices into channels of 8-mers and 11-mers, respectively. Furthermore, molecular dynamics simulations suggest that lipids can favorably assemble into a bilayer within the larger CALHM2 pore, but not within CALHM1, demonstrating the potential correlation between pore-size, lipid accommodation, and channel activity.

## Main Text

First identified as a genetic risk factor for Alzheimer’s disease(*1*) and later as voltage-gated channels expressed in the brain(*2*) and taste cells(*3–6*), CALHM1 has been the most well studied family member to date. The function of CALHM2 proteins expressed in astrocytes have been linked to depression(*7*) and implicated to play a role in glial-neuronal functions(*7*). While CALHM3 has been shown to form heteromeric channels with CALHM1(*8*), the functions of the remaining members, CALHM4-6, are currently unknown. The *calhm* genes are conserved throughout vertebrates and non-vertebrates. Furthermore, CALHM1 from *Caenorhabditis elegans* has been shown to possess similar functional properties to that of human CALHM1 (hCALHM1)(*9*), demonstrating functional conservation throughout diverse species as well.

In this study, we focused on the two major family members, CALHM1 and CALHM2, which are involved in controlling excitability of neurons(*2, 7*). We first conducted expression screening of CALHM orthologues using fluorescence coupled size-exclusion chromatography (FSEC)(*10*), which concluded that chicken CALHM1 (chCALHM1) and human CALHM2 (hCALHM2) show protein size homogeneity suitable for structural analysis. chCALHM1 and human CALHM1 (hCALHM1) have 67.7%/80.2% and 81.4%/93.8% sequence identity/similarity overall and within the transmembrane domains (TMDs), respectively (**fig. S1**). chCALHM1 also contains Asp120, the equivalent residue of which in hCALHM1 (Asp121) has been shown to be critical for its ion channel activity(*11*) (**fig. S1**). Indeed, our patch-clamp electrophysiology shows that chCALHM1 has similar functional properties to hCALHM1 including voltage-sensitivity, calcium sensitive inhibition, and channel blockade by ruthenium red (**Fig. 1A-B**). Both chCALHM1 and hCALHM2 proteins were recombinantly expressed in *Sf*9 insect cells, infected with the respective recombinant baculoviruses(*12*), purified, and reconstituted into lipid nanodiscs prior to vitrification for the cryo-EM study (**fig. S2;** also see Supplementary Methods).

**Fig. 1.**
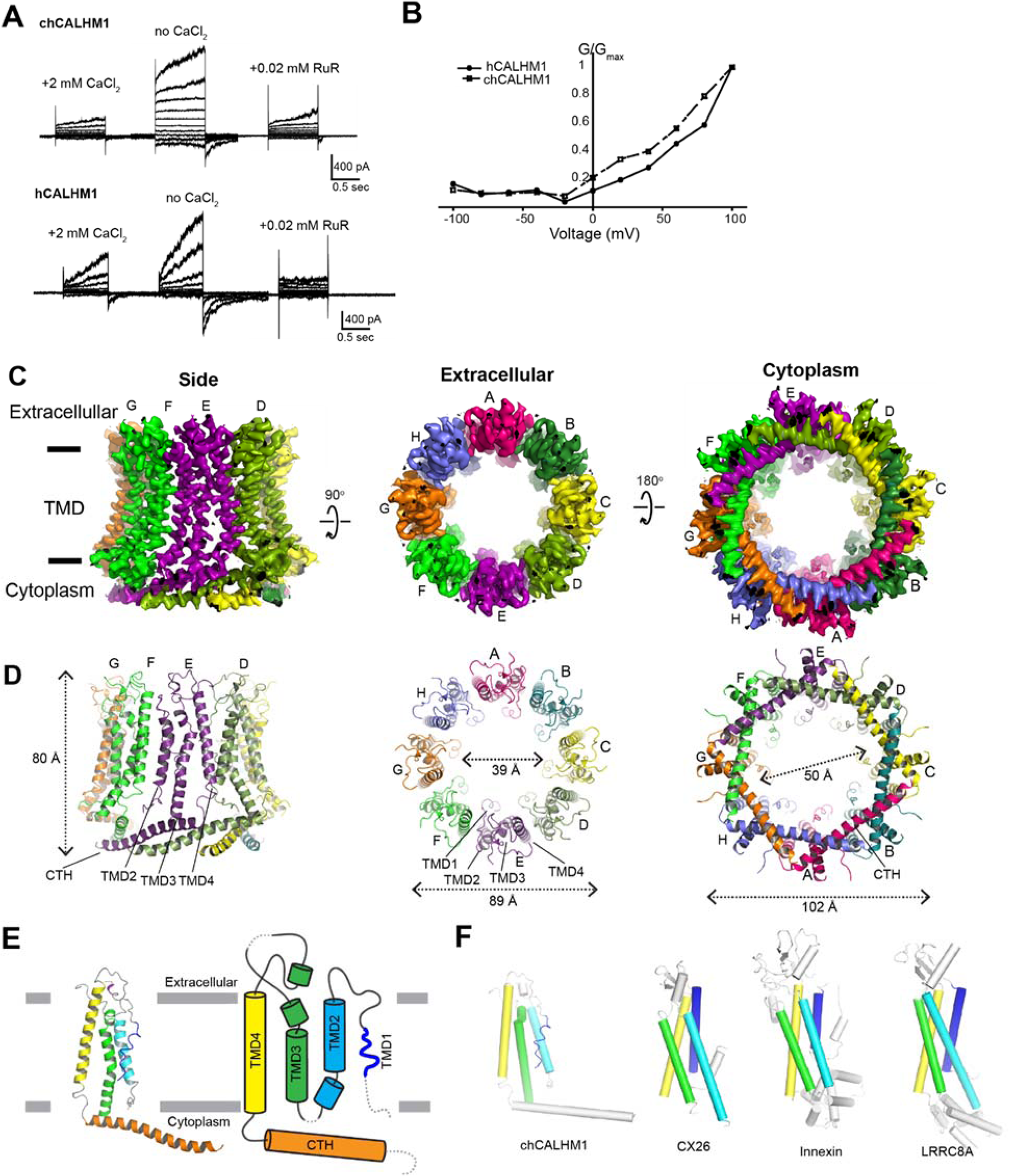
Structure and function of chCALHM1. **(A)** Currents elicited by hCALHM1 and chCALHM1 with voltage steps from −100 to +100 mV in 10 mV increment in the presence and absence of extracellular calcium. The current of chCALHM1 can be blocked by 0.02 mM ruthenium red (RuR) as previously shown for hCALHM1. **(B)** G-V plot of chCALHM1 and hCALHM1 with no CaCl_2_ (*right panel*). **(C-D)** Cryo-EM density (**C**) and atomic models (**D**) of chCALHM1 viewed from the side of the membrane, the extracellular region, and the cytoplasm.**(E)** Ribbon (left) and schematic (right) representations of the chCALHM1 protomer. The TMDs are colored as blue, cyan, green, and yellow for TMD1, 2, 3, and 4, respectively. Dashed lines represent regions that are not visible in our structure. **(F)** Protomers of chCALHM1, CX26(*21*), innexin(*18*), and LRRC8A(*22*). The TMDs are colored as in *panel E* for comparison.

We solved the structure of chCALHM1 using single particle cryo-EM analysis at an overall resolution of 3.63 Å (**Fig. 1C-E, fig. S3-4, table S1**) as assessed by Fourier Shell Correlation (FSC)(*13, 14*). The cryo-EM structure was solved in the absence of calcium, and therefore likely represents the active state. The cryo-EM density of the extracellular domain, the four TMD helices, and the cytoplasmic helices (CTHs) were of sufficient quality to conduct *de novo* modeling between residue numbers 26-79, 91-137 and 151-247, altogether spanning 198 out of 342 amino acids. Most of the missing density is in the carboxyl terminal region after the CTH where 72 out of 94 residues are predicted to be unstructured by a secondary structure analysis(*15*). Nevertheless, the structure confirms the previous prediction that CALHM1 harbors four transmembrane domains with the amino and carboxyl termini facing the cytoplasm(*16*). The cryo-EM density for TMD1 facing the pore is weaker compared to the other three TMDs, indicating the presence of conformational flexibility. Some unresolved density extends from TMD1 towards the middle of the channel at the cytoplasmic side, likely representing the amino terminal residues in multiple conformations (**fig. S5**). In CALHM1s from human and *Caenorhabditis elegans*, the first nine residues have been shown to alter voltage-sensitivity(*17*), thus, we suggest that these voltage-sensing residues are located in the inner-pore within the membrane spanning region (**fig. S5**). Our current structure clearly showed octameric assembly with the pore-like structure in the middle of the oligomer (**Fig. 1C-D**). The assembly is mediated mainly by interactions between TMD2 and TMD4, between TMD1 and TMD3, and between the forty-residue long CTHs of neighboring subunits (**fig. S6**). The octameric assembly shown in our high resolution cryo-EM structure differs from a previous study suggesting hexameric assembly of CALHM1 based on Blue Native-PAGE and photobleaching of the hCALHM1-EGFP constructs(*16*). Nevertheless, the subunit-interface residues are highly conserved between chCALHM1 and hCALHM1 strongly implying preservation of oligomeric mechanisms (**fig. S1 and S6**, 88.5%/100% identity/similarity over 35 residues in TMDs and CTH). The only other octameric channel reported to date is innexin(*18*), however, it does not share similar features in the pattern of oligomeric assembly with chCALHM1. Furthermore, contrary to a previous suggestion, there is no similarity in the membrane topology, structure, and oligomeric assembly pattern of chCALHM1 to *N*-methyl-D-aspartate (NMDA) receptors(*1*) which contain three TMDs and a re-entrant loop and form hetero-tetramers(*19, 20*). Importantly, the structural comparison of monomers demonstrated that chCALHM1 does not resemble other four transmembrane channel proteins including connexin (CX26)(*21*), innexin (INX6)(*18*), or the volume-regulated anion channel (VRAC; LRRC8)(*22, 23*) (**Fig. 1F**).

A key residue known to modulate CALHM1 ion permeability and calcium sensitivity, Asp120 (Asp121 in hCALHM1)(*11*), is located in TMD3 at the interface with the neighboring subunit (**fig. S6A-C**). Although in chCALHM1, each of the Asp120 residues face the inner pore, they do not appear to participate in pore formation directly as they are distant from each other (∼20 Å apart between the Cαs of the neighboring Asp120 residues). Instead, Asp120 may facilitate inter-subunit interactions which stabilize channel assembly (**fig. S6C**). Mutation of another key residue in hCALHM1 (Pro86Leu) has also previously been shown to be a risk factor for the age of onset of Alzheimer’s disease in selected populations(*1*), and although the equivalent residue in chCALHM1 is Gln85, it is clear that this residue is located in the disordered loop between TMD2 and TMD3. Consistent with this location, which faces the cytoplasm and is not part of the channel-pore (**fig. S6A-B;** asterisks), the Pro86Leu mutation in hCALHM1 has previously been shown not to alter channel activity(*1*). Instead, the mechanisms underlying the association of this mutations with Alzheimer’s disease pathology may involve other factors, such as protein-protein interactions involving this loop and subsequent cell signaling events that regulate the level of amyloid beta(*24*).

CALHM2 has a moderately high sequence similarity to CALHM1 within the predicted TMD domains (35%/56% and 37%/55% between hCALHM1 and hCALHM2 and between chCALHM1 and hCALHM2, respectively (identity%/similarity%); **fig. S7**), yet, unlike CALHM1, does not show voltage-dependent ion channel activities(*8*). Thus, we wondered what structural features of CALHM1 and CALHM2 may be responsible for this functional difference. To permit an extensive comparison, we conducted a structural analysis of the human CALHM2 (hCALHM2) protein by implementing a similar protocol to that used above for chCALHM1 (**fig. S8,** also see **Supplementary Methods**). Single particle cryo-EM analysis of the hCALHM2 sample in the absence of calcium resulted in one major 3D class at 3.48 Å resolution as assessed by FSC (**fig. S9-10, table S1**). As in the case of chCALHM1, the hCALHM2 protomer contains four TMDs and the long CTH that is the signature of the CALHM family (**Fig. 2A-B, D**). The orientations of the four TMD helices in hCALHM2 are also similar to that of chCALHM1 and unrelated to connexin (CX26), innexin, or VRAC (**Fig. 2E**). However, the profound structural difference between chCALHM1 and hCALHM2 is that the oligomeric state of hCALHM2 is 11-mer. As in chCALHM1, the oligomeric assembly of hCALHM2 is mediated by interactions between TMD2 and TMD4, between TMD1 and TMD3, and between the CTHs of the neighboring subunits (**fig. S11**). However, the fundamental difference is in the angle between TMD4 and the CTH, which is controlled by the linker sequence that tethers these two helices together (TMD-CTH linker; **Fig. 2E**). Consequently, the sites of inter-CTH interactions are different between chCALHM1 and hCALHM2. This linker sequence differs between CALHM family members, suggesting that CALHM4-6 may also have distinct oligomeric states (**fig. S6**). It is interesting to note that CALHM1 and CALHM3 have similar linker sequences and are known to form heteromers(*8*). Nevertheless, the 11-mer channel assembly observed in hCALHM2 is unprecedented. Therefore, to validate the physiological relevance of this 11-mer assembly in the membrane environment, we conducted a series of disulfide-based inter-subunit crosslinking experiments (**Fig. 2C**). Based on the structure, we substituted residues in the inter-subunit interface at the extracellular side of TMD2 and TMD4 (Arg52/Tyr182) and at the CTHs (Asn226/Arg240) with cysteines (**Fig. 2C**), subjected the mutants to SDS-PAGE, and detected bands by Western blot in the presence and absence of a reducing agent. In the case of CTH mutants located in the cytoplasm, we facilitated disulfide bond formation by addition of copper phenanthroline to the membrane fraction. In the non-reducing condition, we observed a high molecular weight band around 460 kDa. Monomer bands were exclusively observed in the reducing condition, indicating that the band shift in the non-reducing condition is mediated by disulfide bonding between the engineered cysteines (**Fig. 2C**). The resolution of the Western blot experiment is not sufficiently high to unambiguously assign 11-mer assembly, however, the result shows that the oligomeric assembly observed in the cryo-EM structure is consistent with its physiological state in the membrane.

**Fig. 2.**
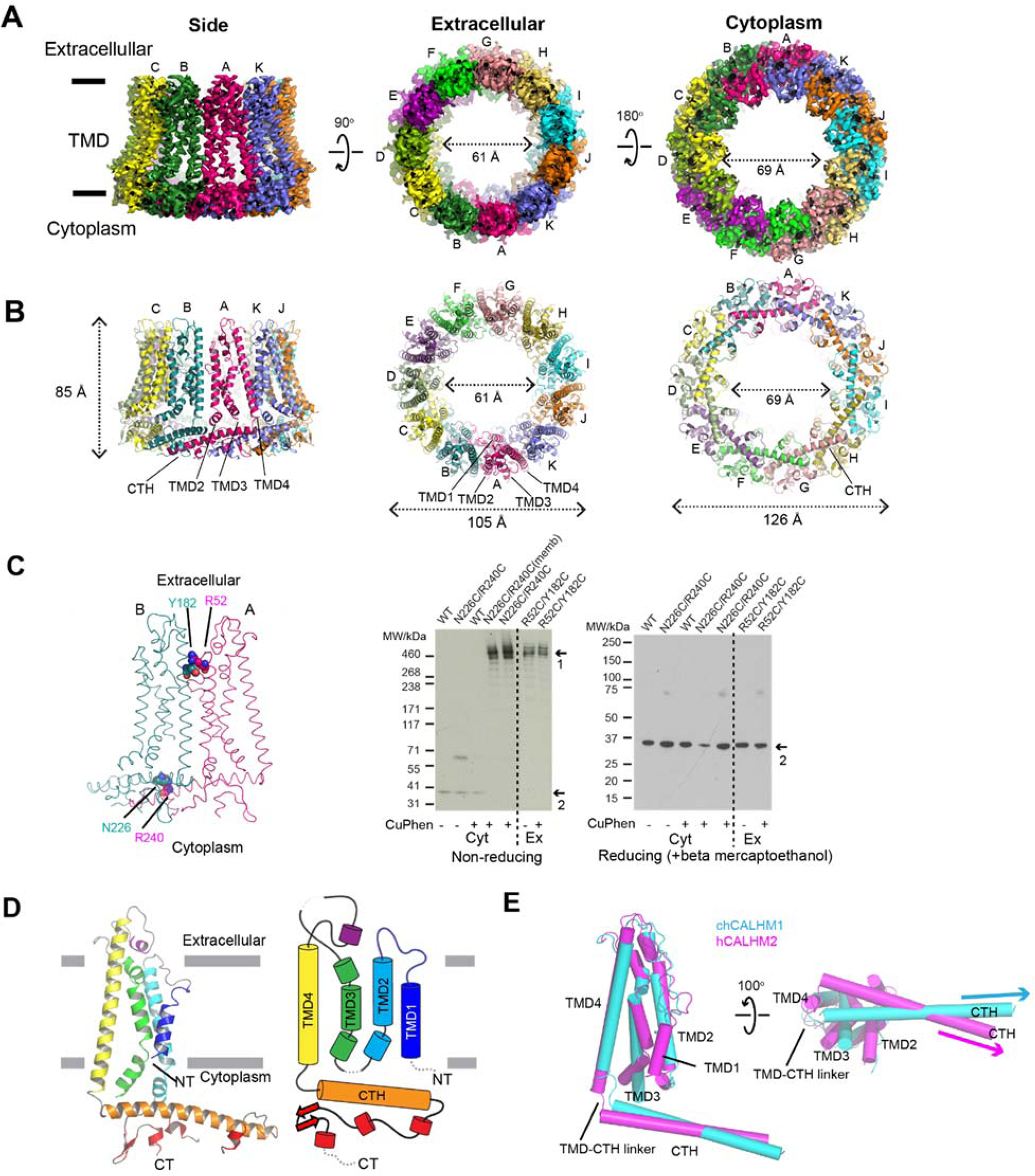
Structure and function of hCALHM2. **(A-B)**, Cryo-EM density (**A**) and atomic models (**B**) of hCALHM2 viewed from the side of the membrane, the extracellular region, and the cytoplasm. **(C)** Cysteine mutations, Tyr182Cys/Arg52Cys or Asn226Cys/Arg240Cys (spheres), were introduced at the subunit interfaces to assess formation of inter-subunit disulfide crosslinking (left). Anti-1D4 Western blots (right) of SDS-PAGE show band shifts for Tyr182Cys/Arg52Cys and Asn226Cys/Arg240Cys (arrow 1). Tyr182Cys/Arg52Cys forms disulfide bonds independent of copper phenanthroline. Formation of disulfide bonds for Asn226Cys/Arg240Cys requires copper phenanthroline treatment before (Asn226Cys/Arg240Cys memb) or after detergent solubilization. The wild-type hCALHM2 protein runs as a monomer under these conditions (arrow 2). Under reducing conditions (beta mercaptoethanol), all constructs run as monomers (arrow 2 on the right gel). **(D)** Ribbon (left) and schematic (right) representations of the hCALHM2 protomer. The TMDs are colored as blue, cyan, green, and yellow for TMD1, 2, 3, and 4, respectively. Dashed lines represent regions that are not visible in our structure. **(E)** Superposition of the TMDs of the chCALHM1 (in cyan) and the hCALHM2 (in magenta) viewed from the side of the membrane (left) and the cytoplasm (right). RMSD of superposition is 1.1 Å over 105 Cα positions.

One noteworthy observation is that the hCALMH2 11-mer dimerizes to form a 22-mer reminiscent of a gap junction in the presence of 1 mM CaCl_2_ under cryo-EM conditions (**fig. S12-13**). Our 22-mer structure solved at 3.68 Å showed a high structural similarity to the 11-mer structure (RMSD = 0.853 Å over 265 residues), indicating that the calcium did not alter the protein architecture. Inclusion of 1 mM CaCl_2_ does not oligomerize hCALHM2 to a 22-mer assembly in solution as observed in the identical peak retention time in size-exclusion chromatography, thus, whether this gap junction structure observed under the cryo-EM condition is physiological or not remains unresolved at this point. However, similar gap junction formation was also observed in cryo-EM images of chCALHM1 under similar conditions (data not shown), indicating the propensity of the CALHM channels to form dimers in the presence of calcium under cryo-EM conditions.

Further structural inspection revealed that the majority of the channel-lining residues (TMD1 and 3) in chCALHM1 are hydrophilic (**Fig. 3A-B**) whereas those in similar positions in hCALHM2 are hydrophobic (**Fig. 3C-E**). Thus, we speculated that hydrophobic molecules, for example lipids, may be favorably placed in this hydrophobic channel-like structure of hCALHM2. This speculation was partially supported by our observation of amorphous density in the middle of the 11-mer assembly in hCALHM2 (**Fig. 3F**), which is not present in the octameric chCALHM1. To assess if the 11-mer channel-like structure in hCALHM2 can accommodate lipids in the middle, we conducted molecular dynamics simulations of chCALHM1 and hCALHM2 in the presence of 1-palmitoyl-2-oleoyl-*sn*-glycero-3-phosphocholine (POPC) (**Fig. 3G-H**). Coarse-grained structural models(*25*) of chCALHM1 and hCALHM2 were mixed with POPC lipids and allowed to self-assemble into bilayers(*26*). During replicates of 500 ns simulations, a membrane bilayer formed around both proteins within the first 100 ns, with a number of lipids inside the channel pore. In hCALHM2, the central pore lipids were oriented with a bilayer-like configuration that remained stable, both in simulations of the protein-membrane systems upon conversion into their corresponding atomistic representation as well as in further, extended coarse-grained simulations (up to 5 µs per replicate) (**Fig. 3H**). By marked contrast, however, although phospholipids could also be accommodated within the smaller chCALHM1 pore, they did not assemble into clearly defined or stable inner and upper leaflets (**Fig. 3H**). These simulations imply that the hydrophobicity and larger 11-mer assembly of hCALHM2 may favor the accommodation of lipid molecules, which prevent ion permeation.

**Fig. 3.**
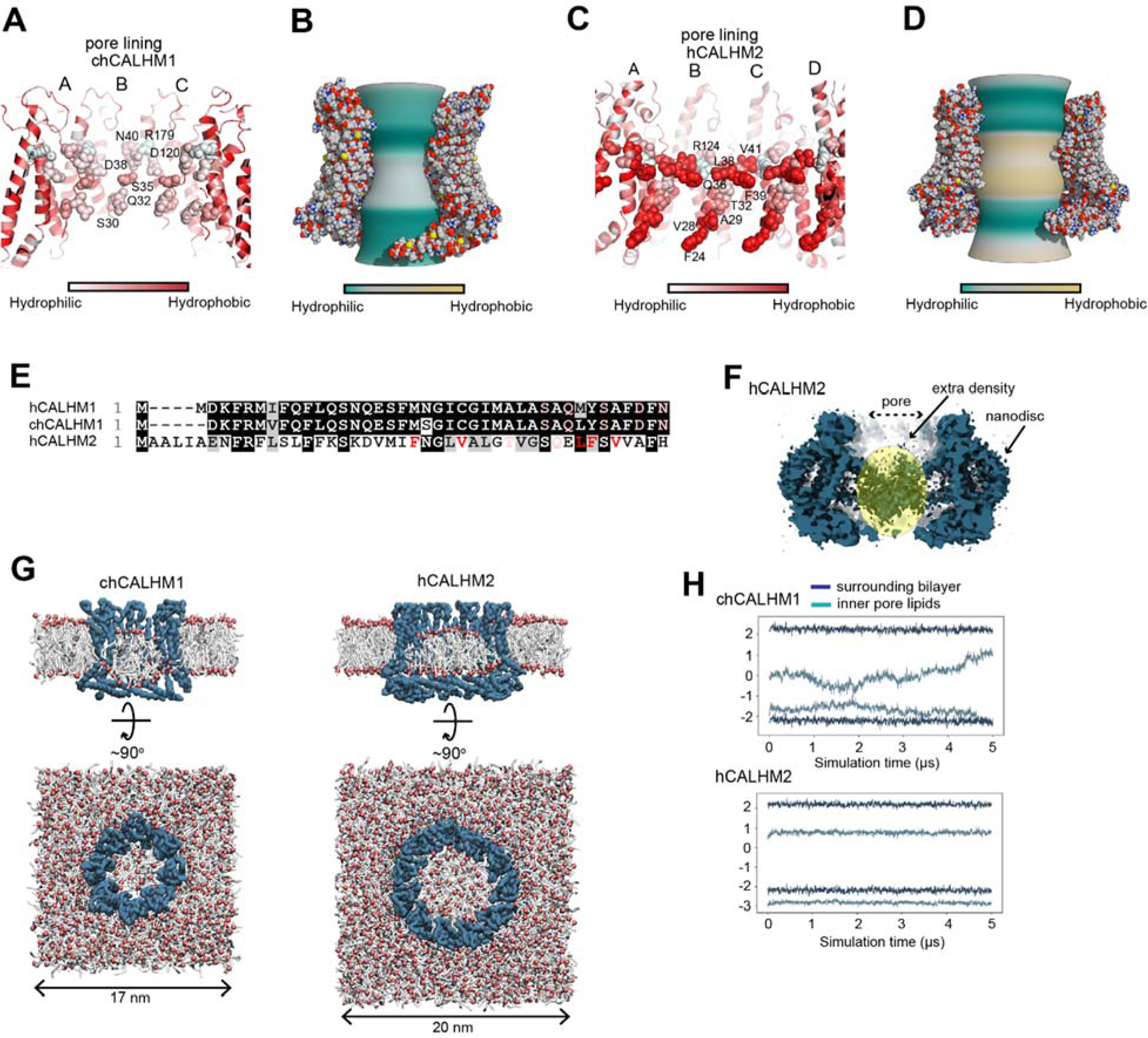
Comparison of pore properties between chCALHM1 and hCALHM2. **(A-D)** Channel lining residues (**A** and **C**)(*27*) and inner pore surface (**B** and **D;** calculated by CHAP(*28*)) of chCALHM1 (**A-B**) and hCALHM2 (**C-D**) colored based on relative hydrophobicity. **(E)** Sequence alignment of the N-terminal residues showing hydrophobic and hydrophilic residues facing the pore with the same color code as in *panel A* and *C*. **(F)** Cross-section of the hCALHM2 showing an extra cryo-EM density in the middle of the pore. **(G)** Coarse-grained MD simulations of chCALHM1 (left) and hCALHM2 (right) embedded in POPC membranes. Side (cutaway) and top views of the final frame of one 5 μs replicate are shown in each case, with the protein backbone particles in blue, phospholipid headgroups in red, and acyl tails in white. Water and ions present in the simulation systems are omitted for clarity. **(H)** Headgroup positions of lipids inside each channel pore and in the surrounding bilayer membrane. The average headgroup z-coordinates of lipids constituting the upper- and lower-leaflets in the final frame of each simulation are respectively tracked through the 5 μs simulated duration; results from one replicate are shown for each protein.

In conclusion, our study demonstrates that CAHLM1 and CALHM2 assemble as 8-mer and 11-mer, respectively, and that these different oligomeric states of CALHMs correlate with channel functions, with only the smaller 8-mer assembly displaying ion channel activity. The structural information presented here will serve as the foundation to study mechanistic questions and reagents development for CALHMs.

## Acknowledgments

We would like to thank D. Thomas and M. Wang for managing the cryo-EM facility and the computing facility at Cold Spring Harbor Laboratory, respectively. A. Hoffmann and T. Malinauskas were involved in the early phase of the research related to hCALHM2.

## Author contributions

J.L.S., .E.C., T.G., N.S., N.G. and H.F. designed and conducted experiments involving cryo-EM. K.M. conducted electrophysiology. S.R. and S.J.T. conducted computational simulations. All authors wrote the manuscript.

## Data and materials availability

Cryo-EM density maps and atomic coordinates of chCALHM1, hCALHM2, and hCALHM2 (22-mer) have been deposited in the electron microscopy data bank under accession codes EMD-####, EMD-####, and EMD-####, respectively.

## Materials and Methods

### Expression, purification, and nanodisc reconstitution of CALHM1 and CALHM2

The chCALHM1 construct with N-terminally fused Strep-II, 8xHis, and EGFP tags (StrepII2-His8-GFP) and the hCALHM2 construct with a C-terminally fused Strep-II tag were expressed in the baculovirus (BV)/*Sf*9 system under the *Drosophila* Hsp70 promoter as previously described(*12*). In brief, *Sf*9 cells were cultured in CCM3 (Invitrogen) supplemented with 1% non-heat inactivated FBS at 27°C, infected with BV at a cell density of 4 × 10^6^ cell/ml, and harvested 48-52 hours after infection. The harvested cell pellets were resuspended in 20 mM Hepes-NaOH (pH 7.5), 200 mM NaCl, 1 mM EDTA and 1 mM PMSF and lysed under high-pressure homogenization (Avestin). The lysate was spun at 4,550*g* for 20 min and the supernatant was ultracentrifuged at 186,000*g* for 1 hour at 4°C. The pellet was solubilized in 20 mM Hepes-NaOH pH 7.5, 200 mM NaCl, 1 mM EDTA, and 1% C12E8 (Anatrace) for 2 hours at 4°C and ultracentrifuged at 186,000*g* for 1 hour at 4°C. The clarified supernatant was loaded onto a Strep-Tactin Sepharose column followed by 20 column volumes (CV) of washing with 20 mM Hepes (pH 7.5), 200 mM NaCl, 1 mM EDTA, 0.01% C12E8 (wash buffer) and elution using the wash buffer supplemented with 3 mM desthiobiotin. The purified hCALHM2 was concentrated to ∼2.5 mg/ml at 4°C using 100-kDa MWCO Amicon concentrators (Millipore) before reconstitution into nanodiscs. Purified chCALHM1 was concentrated to 1 mg/ml, digested by trypsin at a weight-to-weight ratio of 1:20 for 1 hour at 18°C to remove StrepII2-His8-GFP, and purified further by size exclusion chromatography using a Superose 6 10/300 column (GE Healthcare) in 20 mM Tris-HCl (pH 8.0), 200 mM NaCl, 1 mM EDTA, 0.01% C12E8. Peak fractions were pooled and concentrated prior to reconstitution into nanodiscs. For reconstitution into nanodiscs, soybean polar extract, MSP2N2, and the purified CALHM proteins, at final concentrations of 0.75, 0.3 and 0.3 mg/ml, respectively, were mixed for 1 hour at 4°C, followed by detergent removal by SM2 Bio-Beads (BioRad) overnight (∼12 hours). The beads were removed and the solution was further purified by size exclusion chromatography using a Superose 6 10/300 column (GE Healthcare) in 20 mM Tris-HCl pH 8.0, 200 mM NaCl, 1 mM EDTA. Peak fractions were pooled and concentrated to ∼2.5 mg/ml (hCALHM2) or ∼0.6 mg/ml (chCALHM1) for cryo-EM grid preparation. MSP2N2 protein was expressed and purified as previously described(*29*).

### Cryo-EM sample preparation, image collection and single particle analysis

3-4 µl of the CALHM-nanodisc complex was applied to glow-discharged 1.2/1.3 400 mesh C-flat carbon coated copper grids (Protochips). The grids were blotted for 4 s with blot force 7 at 85% humidity and 15°C prior to plunge freezing into liquid ethane using a Vitrobot Mark IV (Thermo Fisher). Datasets were collected using a Titan Krios operated at an acceleration voltage of 300 keV and the GATAN K2 Summit direct electron detector coupled with the GIF quantum energy filter (Gatan Inc.) controlled by SerialEM software(*30*). Movies were recorded with a pixel size of 1.06 Å, an exposure time of 10 s over 50 frames, and a dose rate of 1.4 e/Å^2^/frame. The program Warp was used to align movies, estimate the CTF and pick particles(*31*). 2D classification, *ab-initio* 3D map generation, 3D refinement, 3D classification, per particle CTF refinement and B-factor sharpening were performed using the program cisTEM(*32*). The highest resolution of 3D refinement used was 6 Å for all of the models in this study. The workflows of single particle analyses for chCALHM1 and hCALHM2 are outlined in **fig. S3 and S9**. *De novo* modeling was done manually using the program Coot(*33*). The final models were refined against the cryo-EM maps using PHENIX real space refinement(*34*) with secondary structure and Ramachandran restraints. The FSCs were calculated by phenix.mtriage. Data collection and refinement statistics are summarized in **table S1**.

### Cysteine crosslinking and Western blot analysis

The cysteine substituted Asn226Cys/Arg240Cys hCALHM2 and Arg52Cys/Tyr182Cys hCALHM2, as well as wild-type hCALHM2, were all C-terminally 1D4 tagged and expressed in the BV/*Sf*9 expression system under the CMV promoter. The membrane fraction were isolated and solubilized as above. The solubilized fraction was subjected to affinity purification by 1D4 antibody conjugated to CNBr-activated agarose (GE Healthcare). The resin was extensively washed and the protein eluted with wash buffer supplemented with 0.2 mg/ml 1D4 peptide. Samples were then either reduced with β-mercaptoethanol, left untreated, or treated with the oxidizing agent (1,10-phenanthroline) copper(II). After 30 min incubation on ice, 1 mM final concentration iodoacetamide was added to samples treated with (1,10-phenanthroline) copper(II). Samples were subjected to Western blot using anti-1D4 monoclonal antibodies (University of British Columbia) and anti-mouse Horseradish peroxidase-conjugated antibodies (GE Healthcare). Protein bands were detected by enhanced chemiluminescence on X-ray film (ECL kit; GE Healthcare). To verify that the in-membrane hCALHM2 assembly corresponded in size to detergent extracted hCALHM2, the membrane fractions containing Asn226Cys/Arg240Cys hCALHM2-1D4 were also oxidized with (1,10-phenanthroline) copper(II) prior to detergent solubilization.

### Molecular dynamics

Molecular structures of chCALHM1 and hCALHM2 were separately embedded within POPC bilayer membranes that were solvated on either side at 0.2 M NaCl concentration. Simulation cells were of approximate dimensions 17 × 17 × 15 nm^3^ (CALHM1) and 20 × 20 × 15 nm^3^ (CALHM2). Each protein-membrane system was assembled and equilibrated via a previously established protocol(*26*). Simulations were performed with GROMACS 5.1(*35*). The MARTINI 2.2 force field(*25*) was used for coarse-grained simulations, with a time-step of 20 fs and an elastic network used to harmonically restrain Cα particles and stabilize the protein structure. Atomistic simulations were run using the OPLS all-atom protein force field with united-atom lipids(*36*) and the TIP4P/2005 water model(*37*). The integration time-step was 2 fs. Temperature and pressure were maintained at 37°C and 1 bar during simulations, using the velocity-rescaling thermostat(*38*) in combination with a semi-isotropic Parrinello and Rahman barostat(*39*), with coupling constants of τ_T_ = 0.1 ps and τ_P_ = 1 ps, respectively. Bonds were constrained through the LINCS algorithm(*40*). A Verlet cut-off scheme was applied, and long-range electrostatic interactions were calculated using the Particle Mesh Ewald method(*41*).

### Electrophysiology

CALHM proteins were expressed in HEK293T cells infected by the recombinant BV harboring chCALHM1 or hCALHM under the CMV promoter. Recordings were obtained ∼48 h post infection using borosilicate glass pipettes (Sutter Instruments) pulled and polished to a final resistance of 2-6 MΩ and backfilled with (in mM) 147 NaCl, 10 EGTA, and 10 HEPES pH 7.0 with NaOH. The bath solution contained (in mM) 147 NaCl, 13 glucose, 10 HEPES pH 7.3 with NaOH, 2 KCl, 2 CaCl_2_, and 1 MgCl_2_. Recordings performed in the absence of Ca^2+^ used a similar solution but with no CaCl_2_ added. A rapid solution exchanger (RSC-200; Bio-logic) was used to perfuse cells with various solutions. Data was collected on an AxoPatch 200B patch-clamp amplifier (Axon Instruments), filtered at 2 kHz (Frequency Devices), and digitized with a Digidata 1550B digitizer (Axon Instruments) using a sampling frequency of 10 kHz. Recordings were analyzed using the Clampex 11.0 software (Axon Instruments). Patches were held at −60 mV and stepped between −100 mV and +100 mV in 20 mV increments for 1 s.

---

**Fig. S1.**
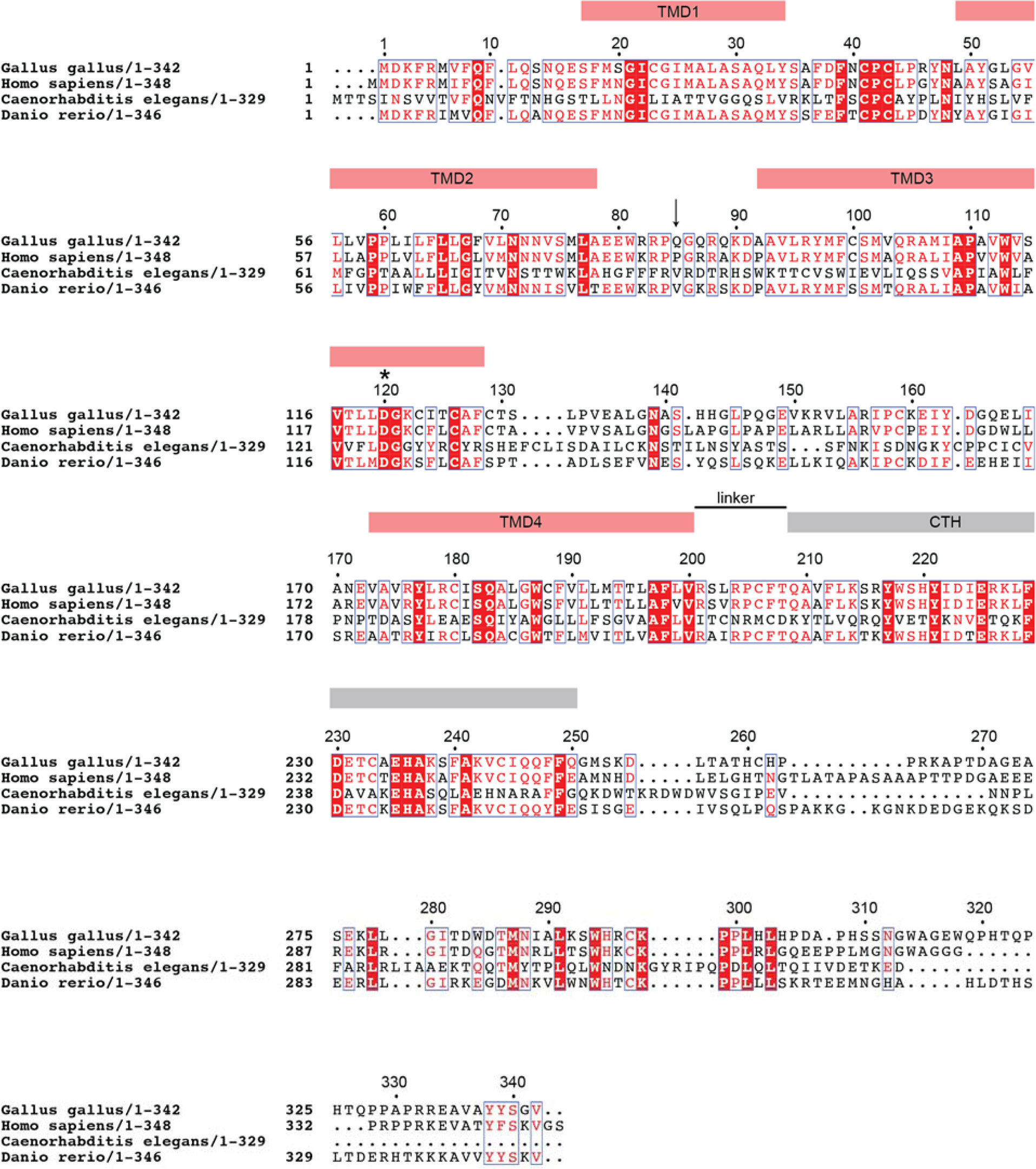
Sequence alignment of CALHM1 orthologues. A multiple sequence alignment of CALHM1 orthologues (*Gallus gallus, Homo sapiens, Caenorhabditis elegans and Danio rerio*). Red boxes indicate identical residues and red characters indicate similar residues. The positions of the TMD1-4 (red bars above the alignment), the CTH (the grey bar above the alignment), and the ‘linker’ are based on the chCALHM1 structure from the current study. An asterisk and an arrow annotate Asp120 and the position of Pro86 in *Homo sapiens* CALHM1 (Glu85 in chCALHM1), respectively. The multiple sequence alignment was generated using Clustal Omega(*42*) and graphically presented using ESPript 3.0(*43*).

**Fig. S2.**
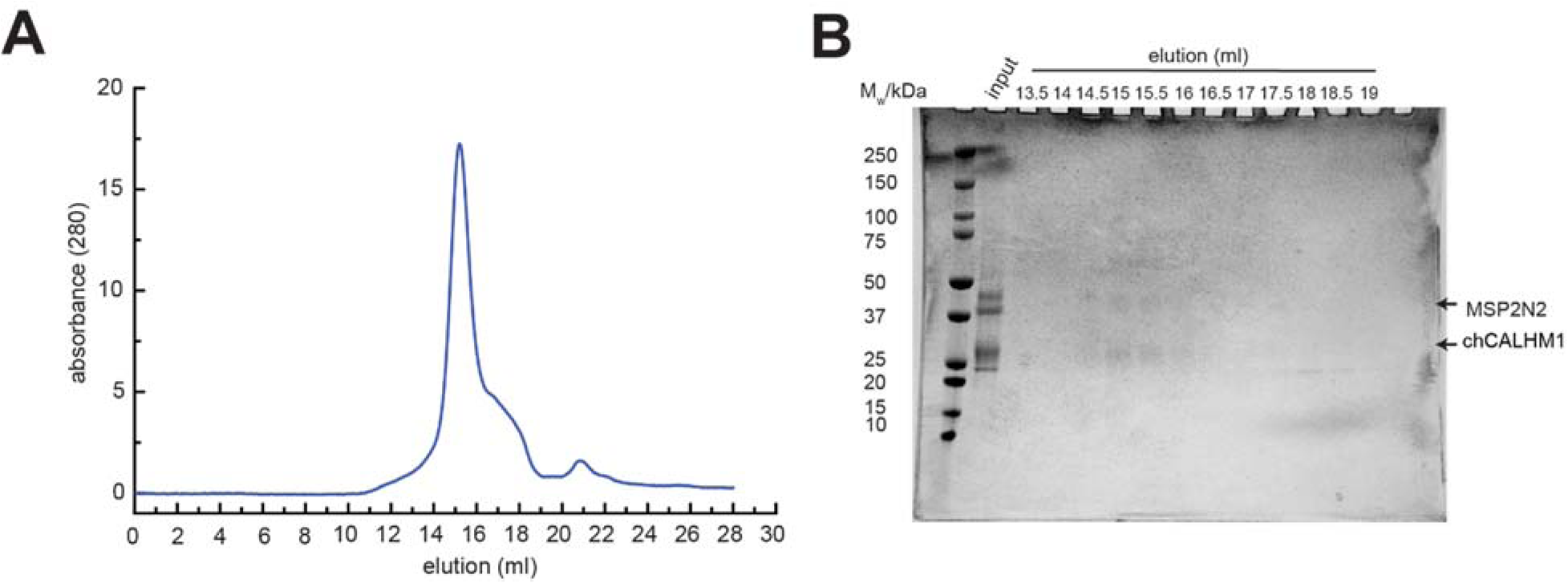
Reconstitution of chCALHM1 into lipid nanodiscs. **(A)** Representative Superose-6 SEC chromatograph of chCALHM1 in MSP2N2 nanodiscs with soy polar extract. **(B)** SDS-PAGE of the fractions collected from SEC. The band for chCALHM1 has a tendency to spread out in SDS-PAGE. Fractions that eluted between 13.5-15.5 ml were pooled, concentrated and subjected to cryo-EM.

**Fig. S3.**
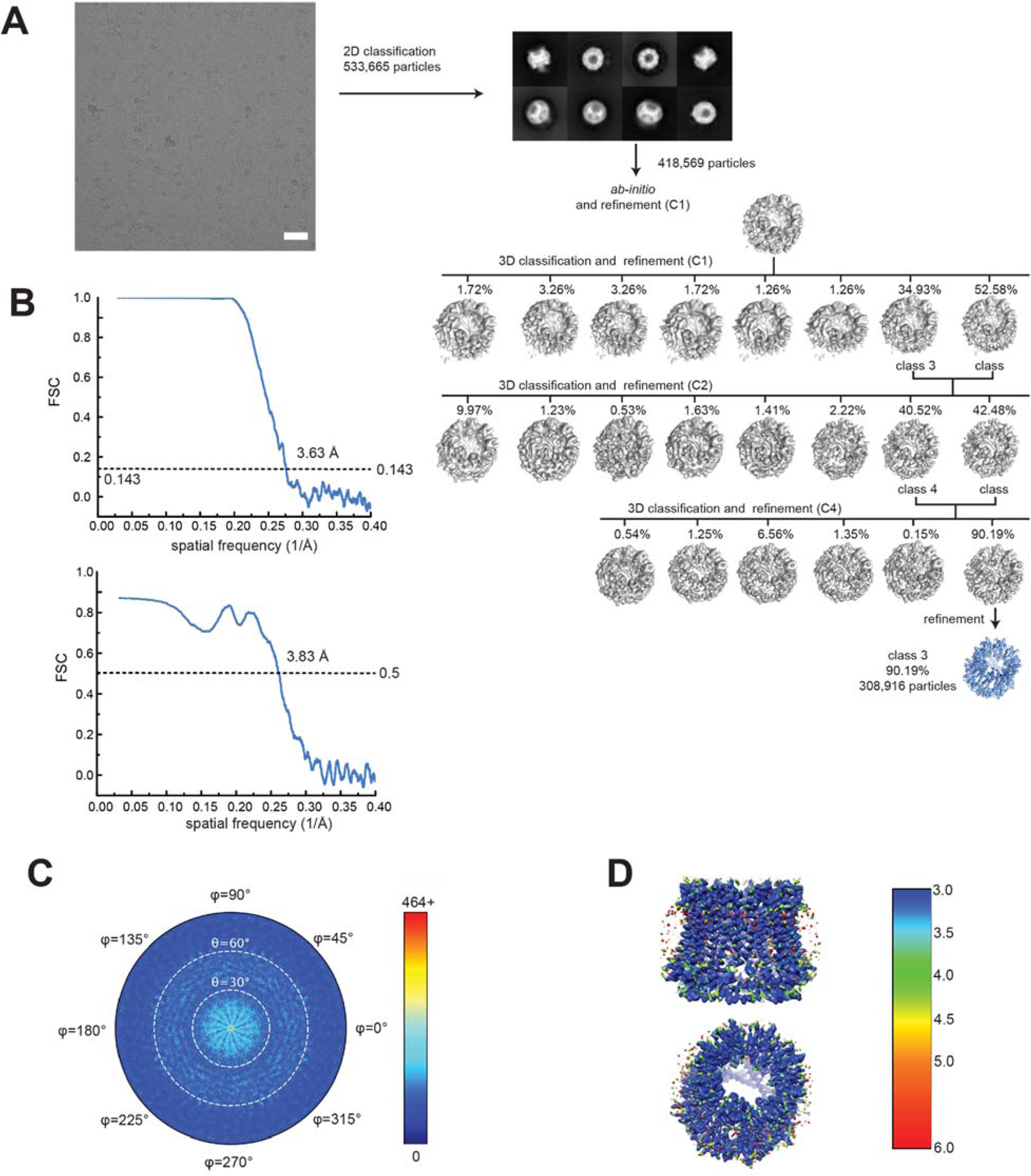
Single particle analysis of chCALHM1. **(A)** A representative micrograph (scale bar = 38.8 nm), representative 2D class averages, and the 3D classification workflow are shown. **(B)** The FSC plots of the two half maps (top) and the map vs model (bottom) are shown. **(C)** The angular distribution plot for class 3. **(D)** Local resolutions of class 3 were calculated using ResMap(*44*).

**Fig. S4.**
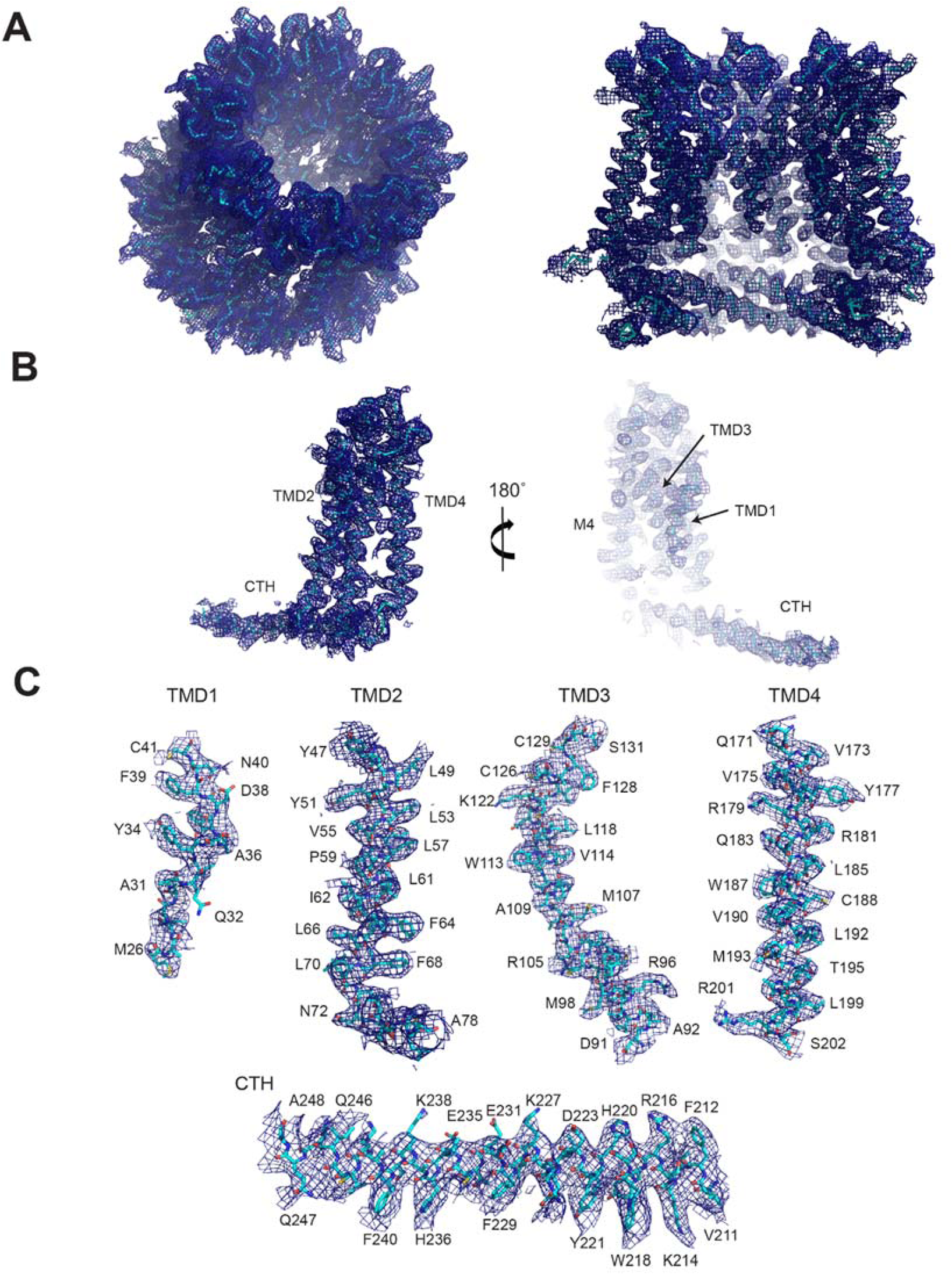
Representative cryo-EM density of chCALHM1. **(A)** Cryo-EM density of the overall octameric assembly (left) and the cross sectional view of the central cavity (right). **(B-C)** Representative density for a monomer **(B)**, and individual TMDs and a CTH **(C)**.

**Fig. S5.**
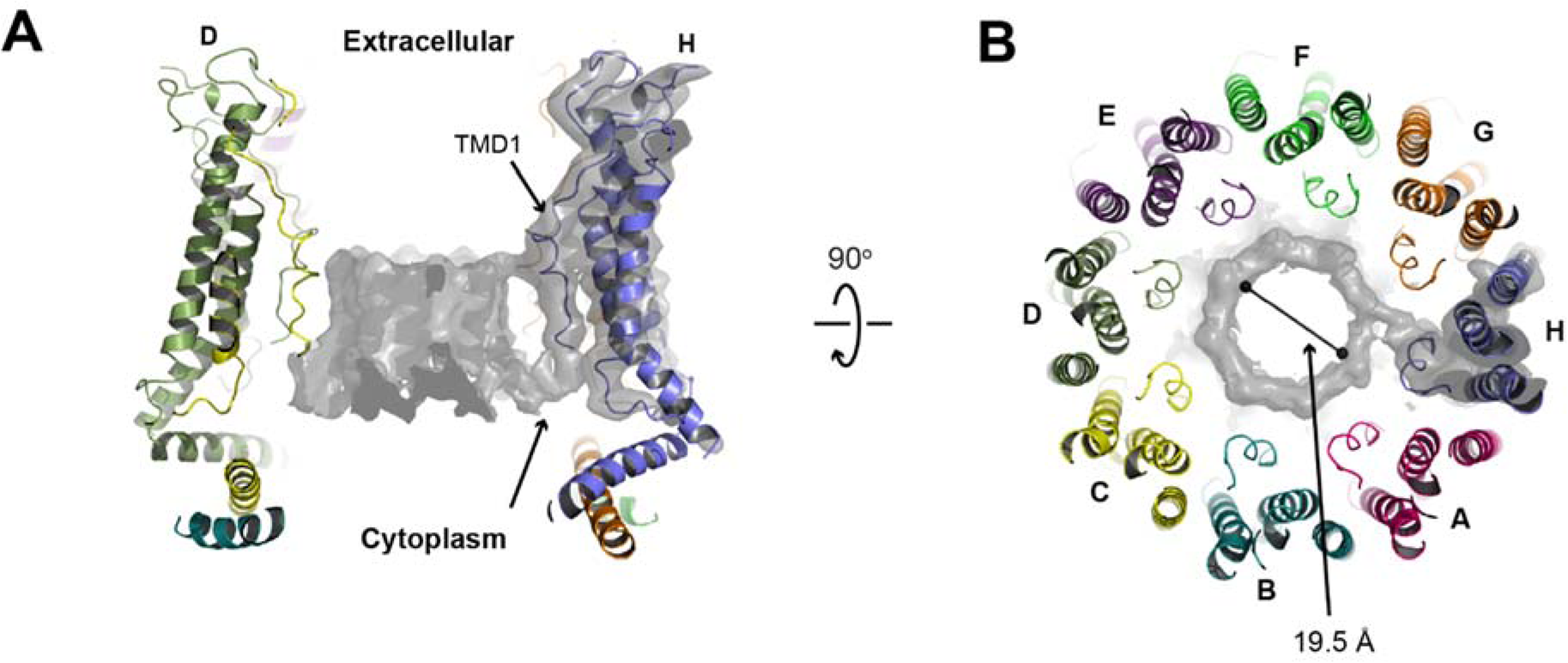
Presence of extra cryo-EM density in the chCALHM1 pore. **(A)** Extra cryo-EM density is observed in the middle of pore-like structure of the chCALHM1. Here the pore-density and the density for only subunit H are shown for clarity. TMD1 and the pore-density are continuous (arrow). **(B)** The density observed from the top of the extracellular region. The diameter of the pore is 19.5 Å.

**Fig. S6.**
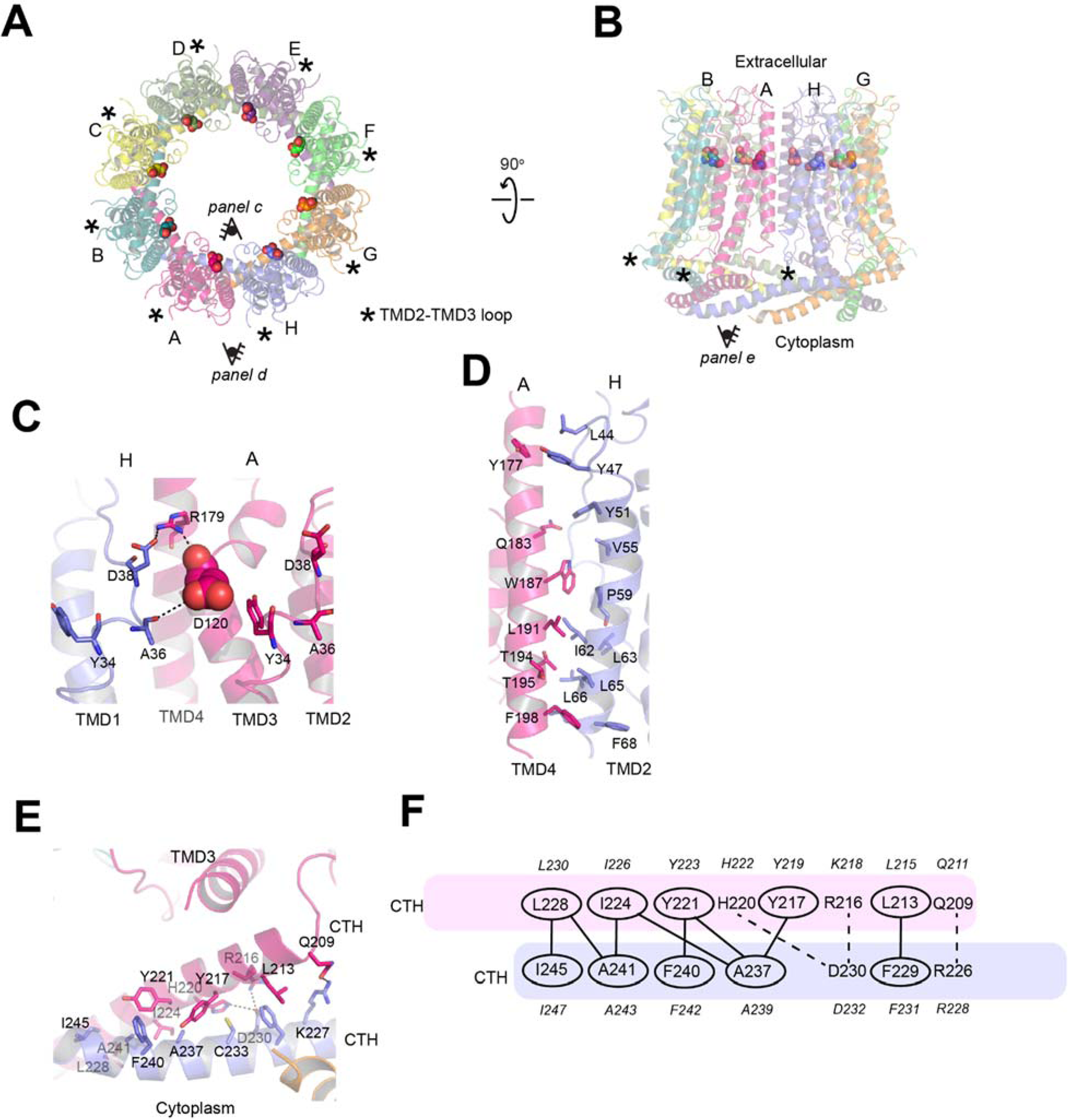
Inter-subunit interface of chCALHM1. **(A-B)** The chCALHM1 structure viewed from the top of extracellular region **(A)** and the side of the membrane **(B)**. Shown in spheres are the Asp120 residues critical for calcium sensitivity and ion permeation. **(C)** Asp120 (sphere) and surrounding residues (sticks) form polar interactions to mediate inter-subunit interactions. **(D-E)** The inter-subunit interactions between TMD2 and TMD4 **(D)** and CTHs **(E)**. **(F)** The schematic presentation of the interactions between two CTHs (magenta and slate blue). Polar and van der Waals interactions mediated by hydrophobic residues (ovals) are shown as dashed and solid lines, respectively. The residues in italic are the equivalent ones in hCALHM1 in a sequence alignment.

**Fig. S7.**
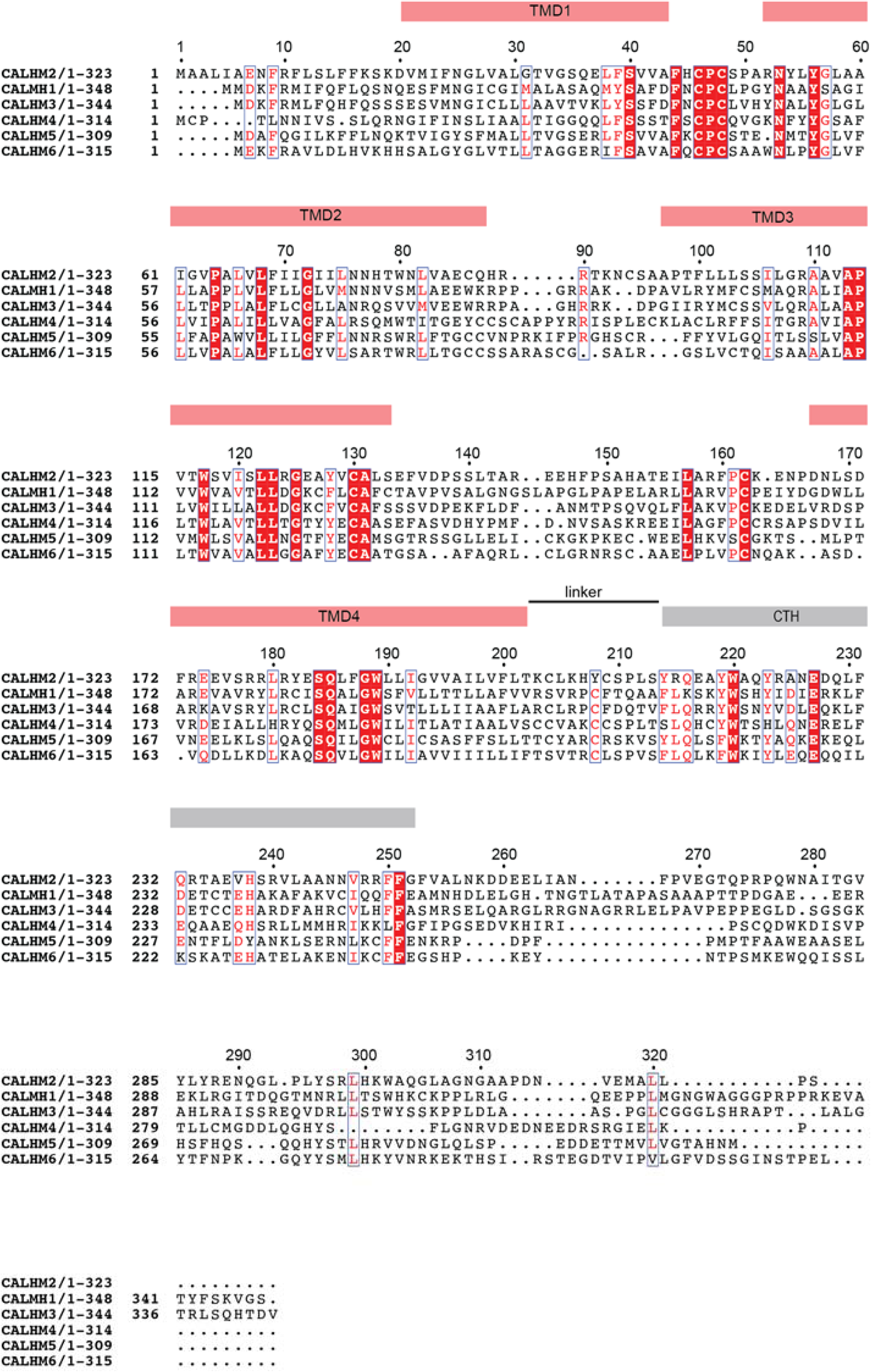
Sequence alignment of CALHM family members. A multiple sequence alignment of the *Homo sapiens* CALHM 1-6. The red boxes indicate identical residues and red characters indicate similar residues. The positions of the TMD1-4 (red bars above the alignment), the CTH (the grey bar above the alignment), and the ‘linker’ are based on the hCALHM2 structure from the current study. The multiple sequence alignment was generated as in fig. S1.

**Fig. S8.**
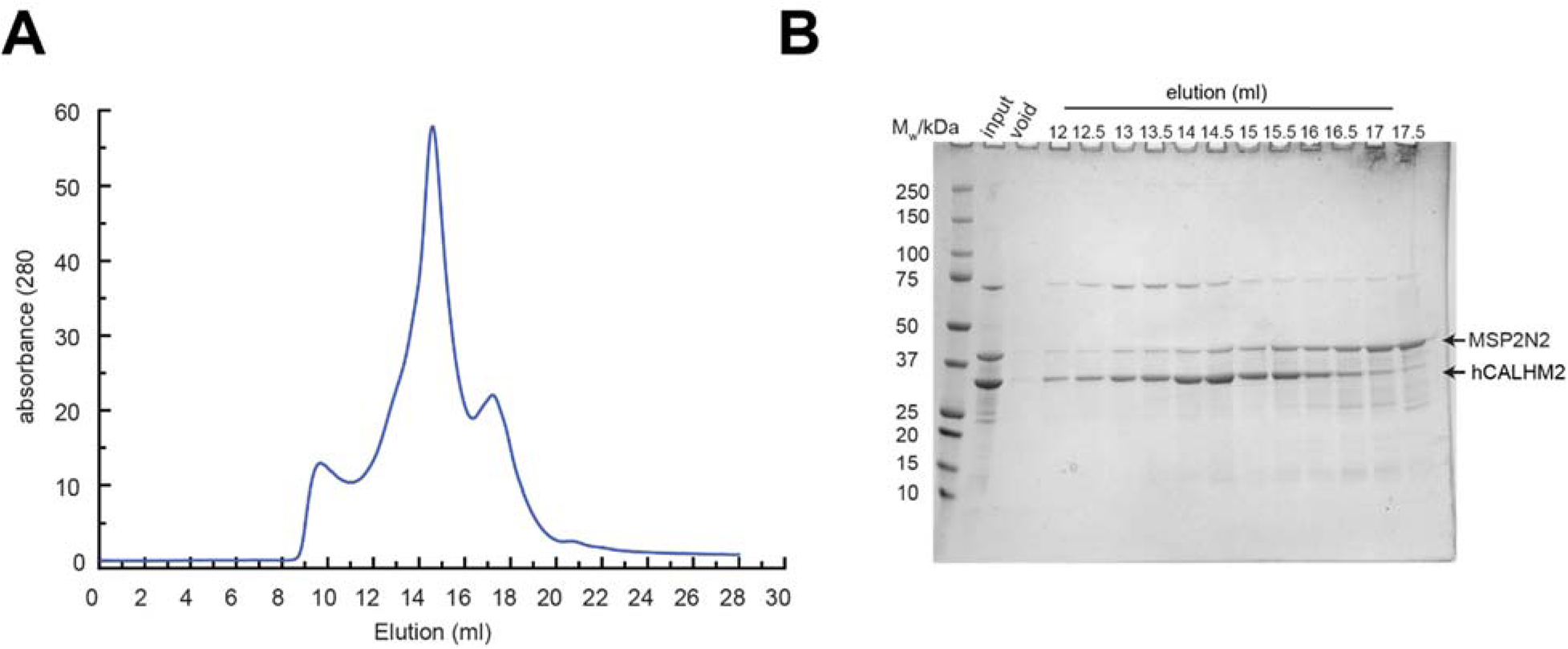
Reconstitution of hCALHM2 into lipid nanodiscs. **(A)** Representative Superose 6 SEC chromatograph of hCALHM2 in MSP2N2 nanodiscs with soy polar extract. **(B)** SDS-PAGE of the fractions collected from SEC. Fractions that eluted between 14.5-16.5 ml were pooled, concentrated and subjected to cryo-EM.

**Fig. S9.**
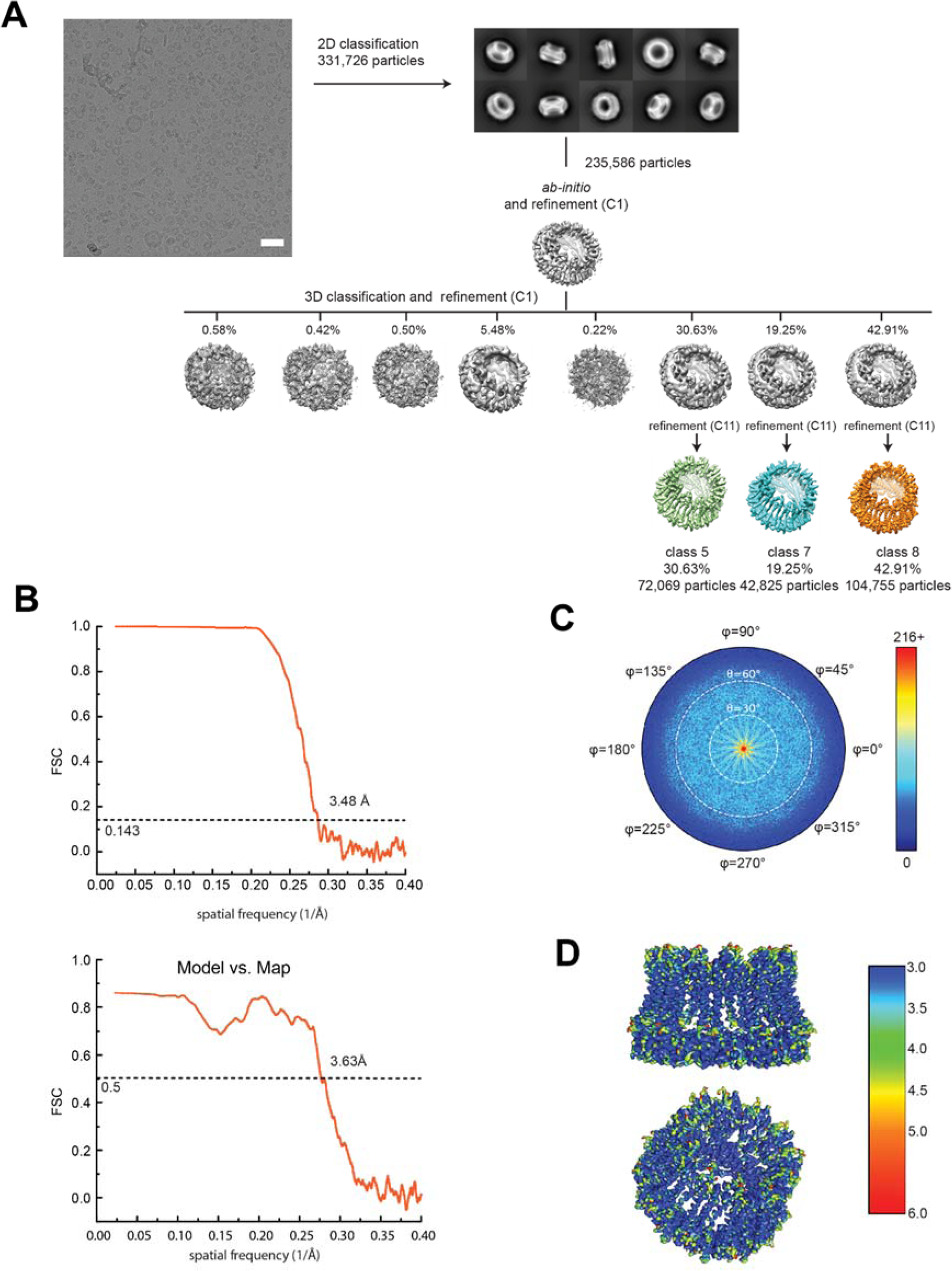
Single particle analysis of hCALHM2. **(A)** A representative micrograph (scale bar = 38.8 nm), representative 2D class averages, and the 3D classification workflow are shown. **(B)** The FSC plots of the two half maps (top) and the map vs. model (bottom) are shown for class 8. **(C)** The angular distribution plot for class 8. **(D)** Local resolutions of class 8 were calculated using ResMap(*44*).

**Fig. S10.**
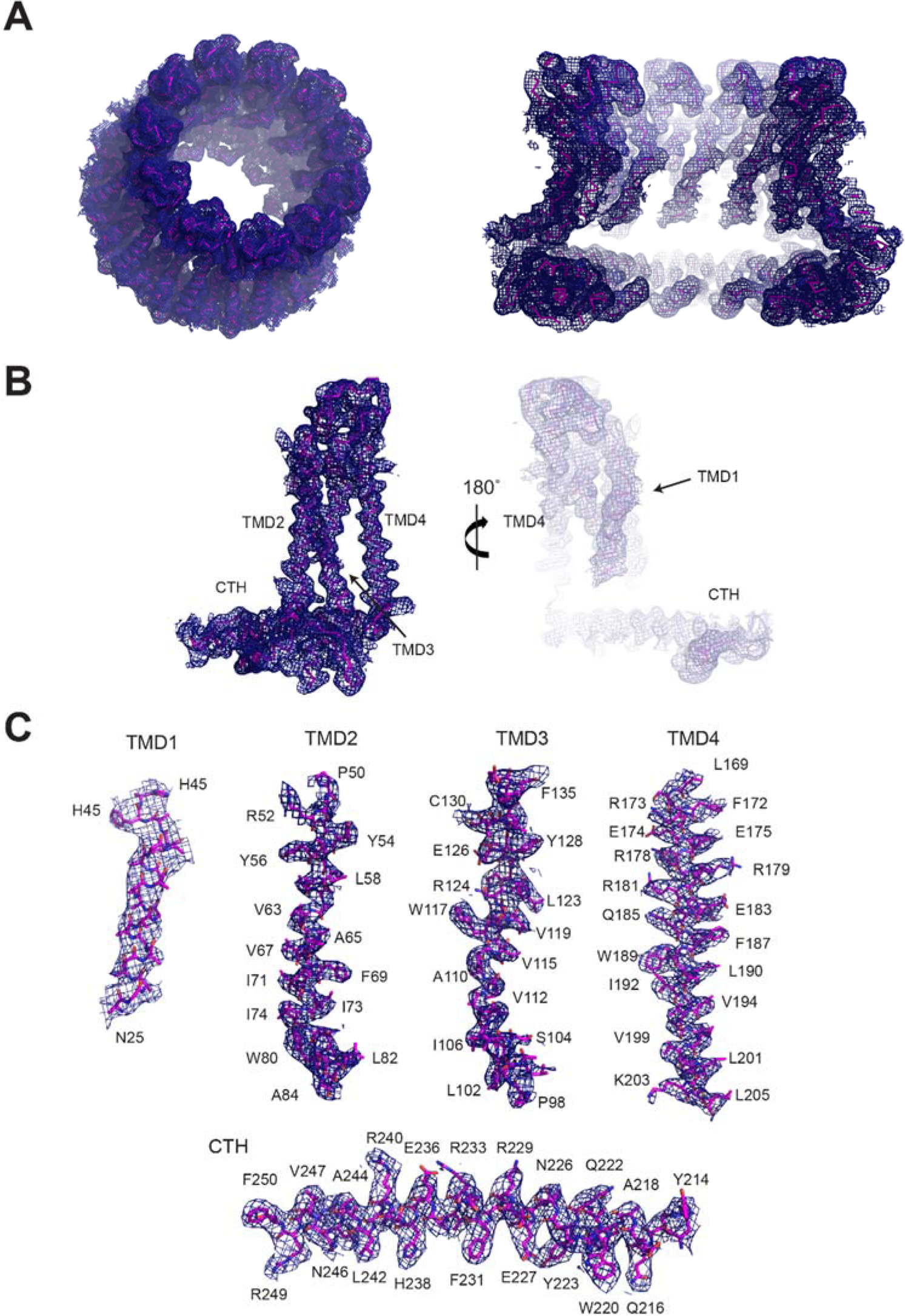
Representative cryo-EM density of hCALHM2. **(A)** Cryo-EM density of the overall 11-mer assembly (left) and the cross-sectional view of the central cavity (right). **(B-C)** Representative density for a monomer **(B)**, and individual TMDs and a CTH **(C)**.

**Fig. S11.**
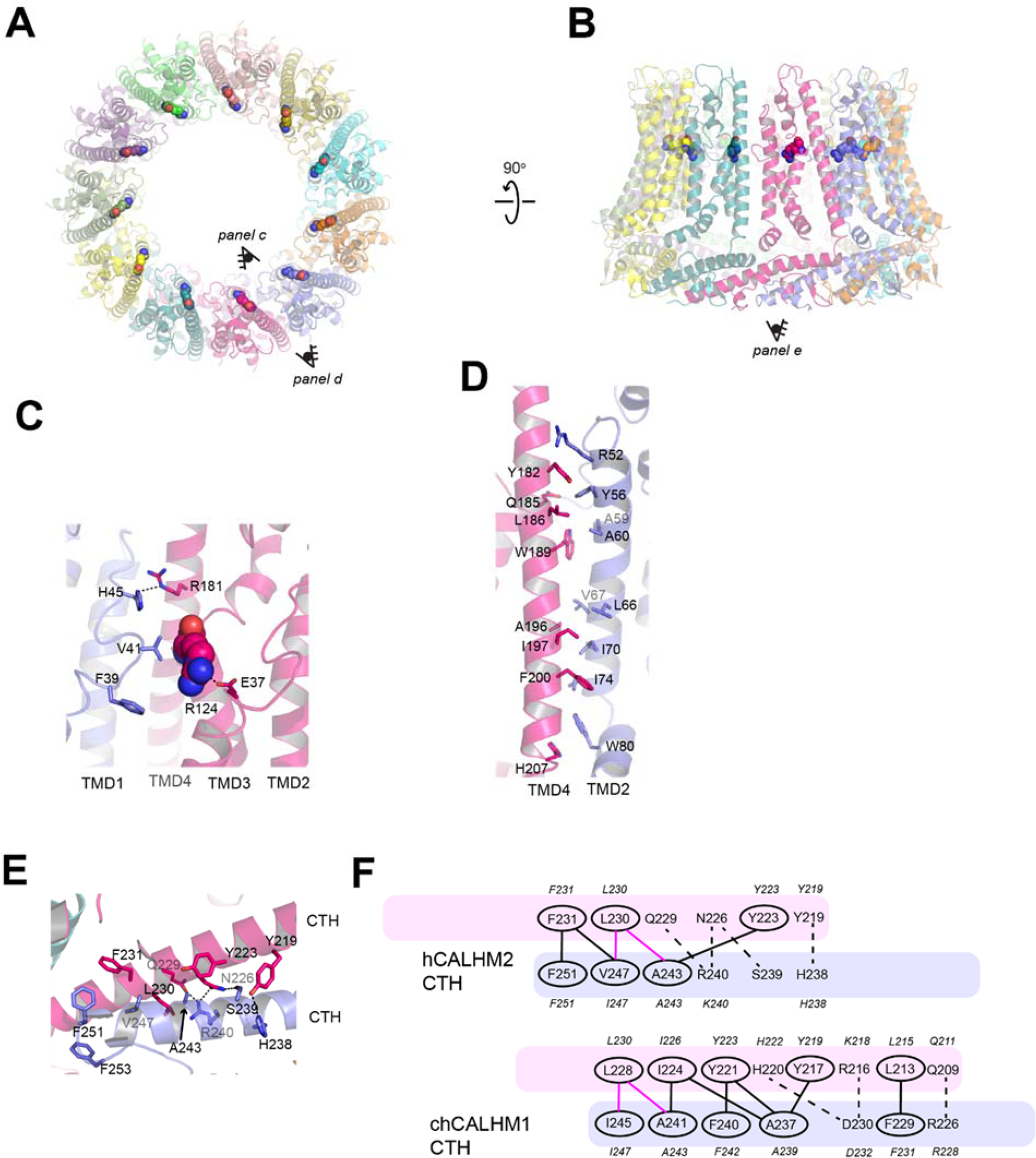
Interaction of hCALHM2 subunits. **(A-B)** The hCALHM2 structure viewed from the top of extracellular region **(A)** and the side of the membrane **(B)**. Shown in spheres are the Arg124 residues at the equivalent position to chCALHM1 Asp120 or hCALHM1 Asp121. **(C)** Arg124 (sphere) and surrounding residues (sticks) form polar and hydrophobic interactions to mediate inter-subunit interactions. **(D-E)** The inter-subunit interactions between TMD2 and TMD4 **(D)** and CTHs **(E)**. **(F)** The schematic presentation of the interactions between two CTHs (magenta and slate blue) in hCALHM2 (top) and chCALHM1 (bottom). Polar and van der Waals interactions mediated by hydrophobic residues (ovals) are shown as dashed and solid lines, respectively. The lines in magenta are the conserved interactions between chCALHM1 and hCALHM2. The residues in italic are the equivalent ones in hCALHM1 in a sequence alignment.

**Fig. S12.**
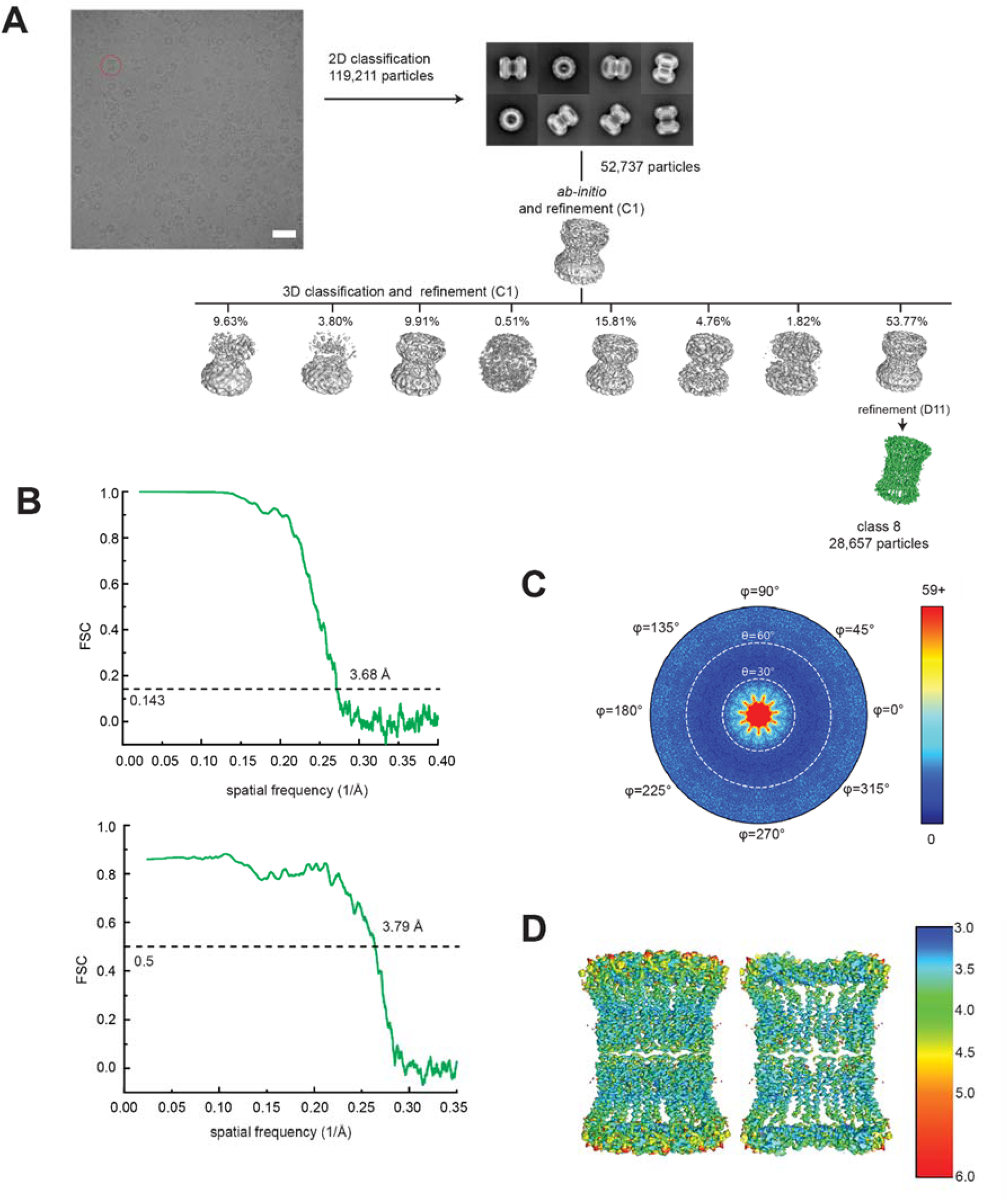
Single particle analysis of 22-meric hCALHM2. **(A)** A representative micrograph (scale bar = 38.8 nm), representative 2D class averages, and the 3D classification workflow are shown. **(B)** The FSC plots of the two half maps (top) and the map vs. model (bottom) are shown for class 8. **(C)** The angular distribution plot for class 8. **(D)** Local resolutions of class 8 were calculated using ResMap(*44*).

**Fig. S13.**
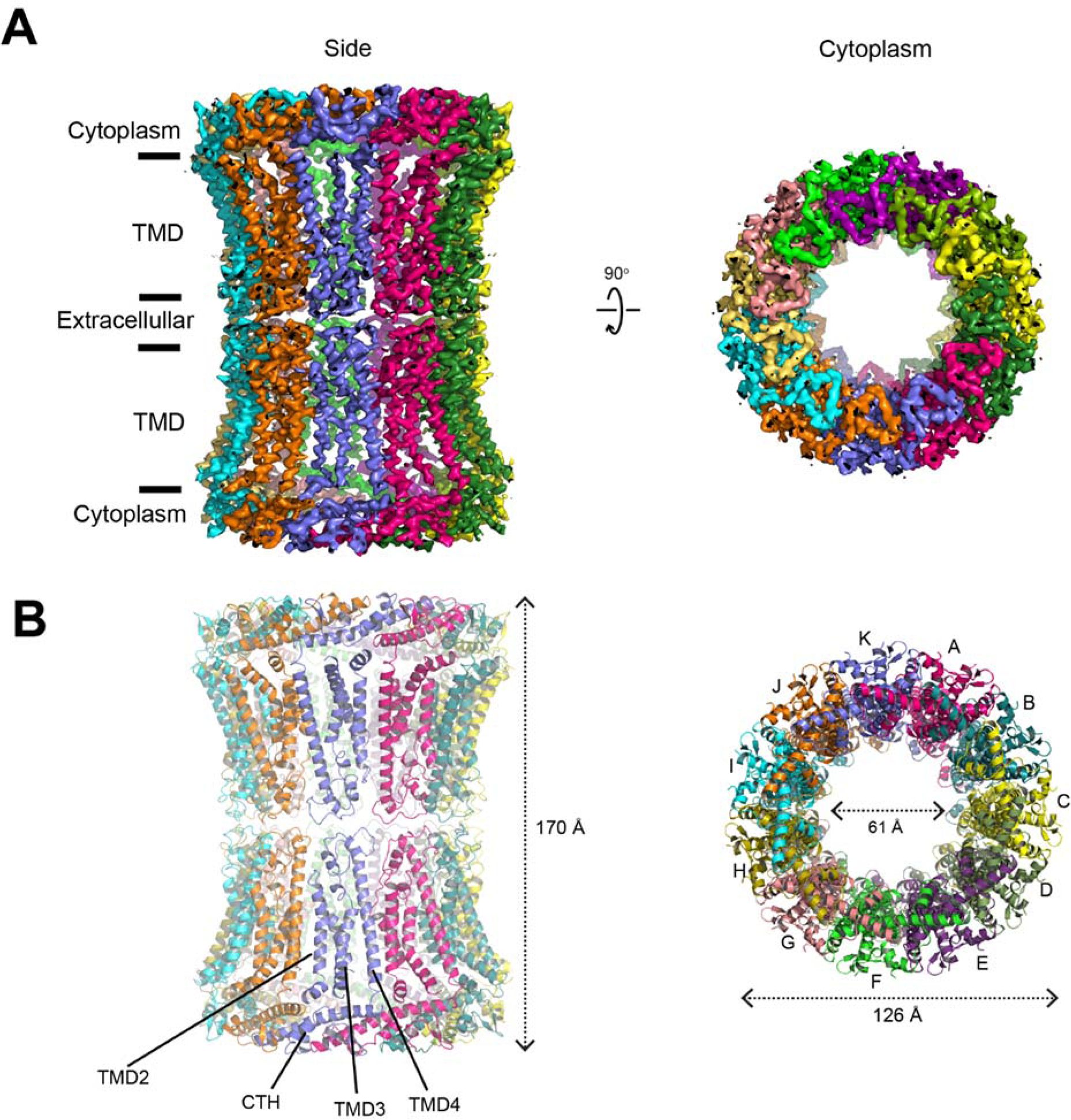
Structure of 22-meric hCALHM2. **(A)** Cryo-EM density of the 22-meric hCALHM2 viewed from the side of the membrane and from the cytoplasm. **(B)** The structural models in the same orientation as the cryo-EM density in **(A)**, showing locations of the TMD2-4 and the CTH. There is little or no structural change between the 22-mer and 11-mer structures except for the extracellular region (due to the inter-11-mer interaction).

**Table S1.** Cryo-EM data collection, refinement and validation statistics.

|  | hCALHM2<br>(EMDB-xxx)<br>(PDB xxxx) | chCALHM1<br>(EMDB-xxxx)<br>(PDB xxxx) | hCALHM2 gap<br>junction<br>(EMDB-xxxx)<br>(PDB xxxx) |
| --- | --- | --- | --- |
| **Data collection and processing** |  |  |  |
| Magnification | 130,000 | 130,000 | 130,000 |
| Voltage (kV) | 300 | 300 | 300 |
| Electron exposure (e-/Å<sup>2</sup>) | 70 | 70 | 70 |
| Defocus range (µm) | 1.2 – 2.8 | 1.5 – 3 | 1.5 – 3 |
| Pixel size (Å) | 1.06 | 1.06 | 1.06 |
| Symmetry imposed | C11 | C8 | D11 |
| Initial particle images (no.) | 331,726 | 533,665 | 119,211 |
| Final particle images (no.) | 104,755 | 308,916 | 52,737 |
| Map resolution (Å) | 3.48 | 3.63 | 3.68 |
| FSC threshold | 0.143 | 0.143 | 0.143 |
| **Refinement** |  |  |  |
| Initial model used (PDB code) | *de novo* | *de novo* | hCALHM2 |
| Model resolution (Å) | 3.63 | 3.83 | 3.79 |
| FSC threshold | 0.5 | 0.5 | 0.5 |
| Model resolution range (Å) |  |  |  |
| Map sharpening *B* factor (Å<sup>2</sup>) | -90 | -150 | -90 |
| Model composition |  |  |  |
| Non-hydrogen atoms | 22,781 | 12,528 | 45,832 |
| Protein residues | 2,292 | 1,592 | 5,916 |
| Ligands | 0 | 0 | 0 |
| CC map vs. model (%) | 0.81 | 0.81 | 0.83 |
| R.m.s. deviations |  |  |  |
| Bond lengths (Å) | 0.004 | 0.006 | 0.007 |
| Bond angles (°) | 0.754 | 0.939 | 0.859 |
| Validation |  |  |  |
| MolProbity score | 2.24 | 2.18 | 2.07 |
| Clashscore | 7.08 | 9.34 | 10.78 |
| Poor rotamers (%) | 2.96 | 0.59 | 0.07 |
| Ramachandran plot |  |  |  |
| Favored (%) | 91.79 | 83.42 | 91.09 |
| Allowed (%) | 8.21 | 15.93 | 8.89 |
| Disallowed (%) | 0 | 0.65 | 0.02 |

## References

1. U. Dreses-Werringloer et al., A polymorphism in CALHM1 influences Ca2+ homeostasis, Abeta levels, and Alzheimer’s disease risk. Cell 133, 1149–1161 (2008).

2. V. Vingtdeux et al., CALHM1 deficiency impairs cerebral neuron activity and memory flexibility in mice. Sci Rep 6, 24250 (2016).

3. A. Taruno et al., CALHM1 ion channel mediates purinergic neurotransmission of sweet, bitter and umami tastes. Nature 495, 223–226 (2013).

4. A. Taruno, I. Matsumoto, Z. Ma, P. Marambaud, J. K. Foskett, How do taste cells lacking synapses mediate neurotransmission? CALHM1, a voltage-gated ATP channel. Bioessays 35, 1111–1118 (2013).

5. Z. Ma, J. E. Tanis, A. Taruno, J. K. Foskett, Calcium homeostasis modulator (CALHM) ion channels. 395–403 (2016).

6. Z. Ma, W. T. Saung, J. K. Foskett, Action Potentials and Ion Conductances in Wild-type and CALHM1-knockout Type II Taste Cells. Journal of Neurophysiology, jn.00835.02016 (2017).

7. M. Jun et al., Calhm2 governs astrocytic ATP releasing in the development of depression-like behaviors. Mol Psychiatry 23, 1091 (2018).

8. Z. Ma et al., CALHM3 Is Essential for Rapid Ion Channel-Mediated Purinergic Neurotransmission of GPCR-Mediated Tastes. Neuron 98, 547–561 e510 (2018).

9. J. E. Tanis et al., Cellular/Molecular CLHM-1 is a Functionally Conserved and Conditionally Toxic Ca 2ϩ -Permeable Ion Channel in Caenorhabditis elegans. J Neurosci, (2013).

10. T. Kawate, E. Gouaux, Fluorescence-Detection Size-Exclusion Chromatography for Precrystallization Screening of Integral Membrane Proteins. Structure 14, 673–681 (2006).

11. Z. Ma et al., Calcium homeostasis modulator 1 (CALHM1) is the pore-forming subunit of an ion channel that mediates extracellular Ca2+ regulation of neuronal excitability. Proc Natl Acad Sci U S A 109, E1963–1971 (2012).

12. M. C. Regan et al., Structural Mechanism of Functional Modulation by Gene Splicing in NMDA Receptors. Neuron 98, 521–529 e523 (2018).

13. G. Harauz, M. van Heel, Exact Filters for General Geometry Three Dimensional Reconstruction. Optik 73, 146–156 (1986).

14. P. B. Rosenthal, R. Henderson, Optimal determination of particle orientation, absolute hand, and contrast loss in single-particle electron cryomicroscopy. J Mol Biol 333, 721–745 (2003).

15. L. Slabinski et al., XtalPred: a web server for prediction of protein crystallizability. Bioinformatics 23, 3403–3405 (2007).

16. A. P. Siebert et al., Structural and functional similarities of calcium homeostasis modulator 1 (CALHM1) ion channel with connexins, pannexins, and innexins. Journal of Biological Chemistry 288, 6140–6153 (2013).

17. J. E. Tanis, Z. M. Ma, J. K. Foskett, The NH2 terminus regulates voltage-dependent gating of CALHM ion channels. Am J Physiol-Cell Ph 313, C173–C186 (2017).

18. A. Oshima et al., Atomic structure of the innexin-6 gap junction channel determined by cryo-EM. Nature Communications 7, 13681 (2016).

19. E. Karakas, H. Furukawa, Crystal structure of a heterotetrameric NMDA receptor ion channel. Science 344, 992–997 (2014).

20. C. H. Lee et al., NMDA receptor structures reveal subunit arrangement and pore architecture. Nature 511, 191–197 (2014).

21. S. Maeda et al., Structure of the connexin 26 gap junction channel at 3.5 A resolution. Nature 458, 597–602 (2009).

22. D. Deneka, M. Sawicka, A. K. M. Lam, C. Paulino, R. Dutzler, Structure of a volume-regulated anion channel of the LRRC8 family. Nature 558, 254–259 (2018).

23. G. Kasuya et al., Cryo-EM structures of the human volume-regulated anion channel LRRC8. Nat Struct Mol Biol 25, 797–804 (2018).

24. V. Vingtdeux et al., CALHM1 ion channel elicits amyloid-beta clearance by insulin-degrading enzyme in cell lines and in vivo in the mouse brain. J Cell Sci 128, 2330–2338 (2015).

25. D. H. de Jong et al., Improved Parameters for the Martini Coarse-Grained Protein Force Field. J Chem Theory Comput 9, 687–697 (2013).

26. P. J. Stansfeld et al., MemProtMD: Automated Insertion of Membrane Protein Structures into Explicit Lipid Membranes. Structure 23, 1350–1361 (2015).

27. D. Eisenberg, E. Schwarz, M. Komaromy, R. Wall, Analysis of membrane and surface protein sequences with the hydrophobic moment plot. Journal of Molecular Biology 179, 125–142 (1984).

28. G. Klesse, S. Rao, M. S. P. Sansom, S. J. Tucker, CHAP: A Versatile Tool for the Structural and Functional Annotation of Ion Channel Pores. J Mol Biol 431, 3353–3365 (2019).

## References

6. Z. Ma, W. T. Saung, J. K. Foskett, Action Potentials and Ion Conductances in Wild-type and CALHM1-knockout Type II Taste Cells. Journal of Neurophysiology jn.00835.02016 (2017).

29. T. K. Ritchie et al., Chapter 11 Reconstitution of Membrane Proteins in Phospholipid Bilayer Nanodiscs. Methods in Enzymology 464, 211–231 (2009).

30. M. Schorb, I. Haberbosch, W. J. H. Hagen, Y. Schwab, D. N. Mastronarde, electron microscopy. Nature Methods.

31. D. Tegunov, P. Cramer, Real-time cryo-electron microscopy data preprocessing with Warp. Nat Methods, (2019).

32. T. Grant, A. Rohou, N. Grigorieff, cisTEM, user-friendly software for single-particle image processing. Elife 7, (2018).

33. P. Emsley, B. Lohkamp, W. G. Scott, K. Cowtan, Features and development of Coot. Acta Crystallogr D Biol Crystallogr 66, 486–501 (2010).

34. P. D. Adams et al., PHENIX: a comprehensive Python-based system for macromolecular structure solution. Acta Crystallogr D Biol Crystallogr 66, 213–221 (2010).

35. M. J. Abraham et al., GROMACS: High performance molecular simulations through multi-level parallelism from laptops to supercomputers. SoftwareX 1-2, 19–25 (2015).

36. W. L. Jorgensen, D. S. Maxwell, J. Tirado-Rives, Development and Testing of the OPLS All-Atom Force Field on Conformational Energetics and Properties of Organic Liquids. Journal of the American Chemical Society 118, 11225–11236 (1996).

37. J. L. F. Abascal, C. Vega, A general purpose model for the condensed phases of water: TIP4P/2005. The Journal of Chemical Physics 123, 234505 (2005).

38. G. Bussi, D. Donadio, M. Parrinello, Canonical sampling through velocity rescaling. The Journal of Chemical Physics 126, 014101 (2007).

39. M. Parrinello, A. Rahman, Polymorphic transitions in single crystals: A new molecular dynamics method. Journal of Applied Physics 52, 7182–7190 (1981).

40. B. Hess, H. Bekker, H. J. C. Berendsen, J. G. E. M. Fraaije, LINCS: A Linear Constraint Solver for molecular simulations. Journal of Computational Chemistry 18, 1463–1472 (1997).

41. T. Darden, D. York, L. Pedersen, Particle mesh Ewald: An N·log(N) method for Ewald sums in large systems. The Journal of Chemical Physics 98, 10089–10092 (1993).

42. F. Madeira et al., The EMBL-EBI search and sequence analysis tools APIs in 2019. Nucleic Acids Research 47, W636–W641 (2019).

43. X. Robert, P. Gouet, Deciphering key features in protein structures with the new ENDscript server. Nucleic Acids Research 42, 320–324 (2014).

44. A. Kucukelbir, F. J. Sigworth, H. D. Tagare, Quantifying the local resolution of cryo-EM density maps. Nature Methods 11, 63–65 (2014).

